# Positive feedback regulation of *Pparγ1sv* and *Pparγ2* during adipogenesis in 3T3-L1 cells

**DOI:** 10.1101/2023.08.03.551916

**Authors:** Yasuhiro Takenaka, Yoshihiko Kakinuma, Ikuo Inoue

## Abstract

We have previously identified the novel splicing variant of mouse *Pparγ* (*Pparγ1sv*) and proposed the synergistic regulation of the early stage of adipocyte differentiation by *Pparγ1sv* and *Pparγ2*. Here, we report the finding of PPARγ-binding sites within the *Pparγ* gene locus and its importance in adipogenesis and propose the positive feedback regulation of *Pparγ1sv* and *Pparγ2* expression during the adipocyte differentiation of 3T3-L1 cells.

## Description

Peroxisome proliferator-activated receptors (PPARs) are nuclear receptors that regulate the expression of several genes involved in metabolic homeostasis. PPARγ, one of the three PPAR gene subtypes, is one of the master regulators of adipogenesis (Tontonoz and Spiegelman, 2008). In a ligand-dependent manner, PPARγ regulates the transcription of target genes, such as *Fabp4* and *C/EBPα*, which are indispensable for the completion of adipocyte differentiation. PPARγ has two isoforms: ubiquitously expressed PPARγ1 and adipocyte-specific PPARγ2. In mice, PPARγ2 is longer than PPARγ1 by 30 amino acid residues at the N-terminus. In addition to *Pparγ1* and *Pparγ2* mRNAs, we previously reported a novel mouse *Pparγ* splicing variant, *Pparγ1sv*, which encodes the PPARγ1 protein and is significantly upregulated during adipocyte differentiation of 3T3-L1 cells and mouse primary cultured preadipocytes (Takenaka et al., 2013). Knockdown of *Pparγ1sv* by specific siRNAs completely abolished the induced adipogenesis in 3T3-L1 cells.

Although the siRNA (siγ1sv22) that we designed (Fig. 1A) specifically knocked-down *Pparγ1sv* transcripts in 3T3L1 cells (Takenaka et al., 2013) (Fig. 1B), we also noticed a notable reduction in *Pparγ2* mRNA levels (Fig. 1C) during adipocyte differentiation of 3T3-L1 cells by quantitative PCR (qPCR) assay. Similarly, siRNA specific to *Pparγ2* (siγ2_8) significantly reduced *Pparγ1sv* levels (Fig. 1B). These results imply that their protein products, PPARγ1 and PPARγ2, mutually affect gene expression. To reveal the presence of cis-acting elements located within or around the *Pparγ* gene locus, we analyzed deposited chromatin immunoprecipitation (ChIP) sequencing study using an anti-PPARγ antibody (Mikkelsen et al., 2010) and found three candidate PPARγ-binding sites (sites 1–3) (Fig. 1D). The binding of CREB-Bindng Protein (CBP) to sites 2–3 and C/EBPα to site 3 supported the notion that these are possible PPARγ-binding sites. We confirmed the binding of PPARγ protein to all three binding sites during addipocyte differentiation by ChIP-qPCR assay using an anti-PPARγ antibody (Fig. 1E).

**Figure 1.**
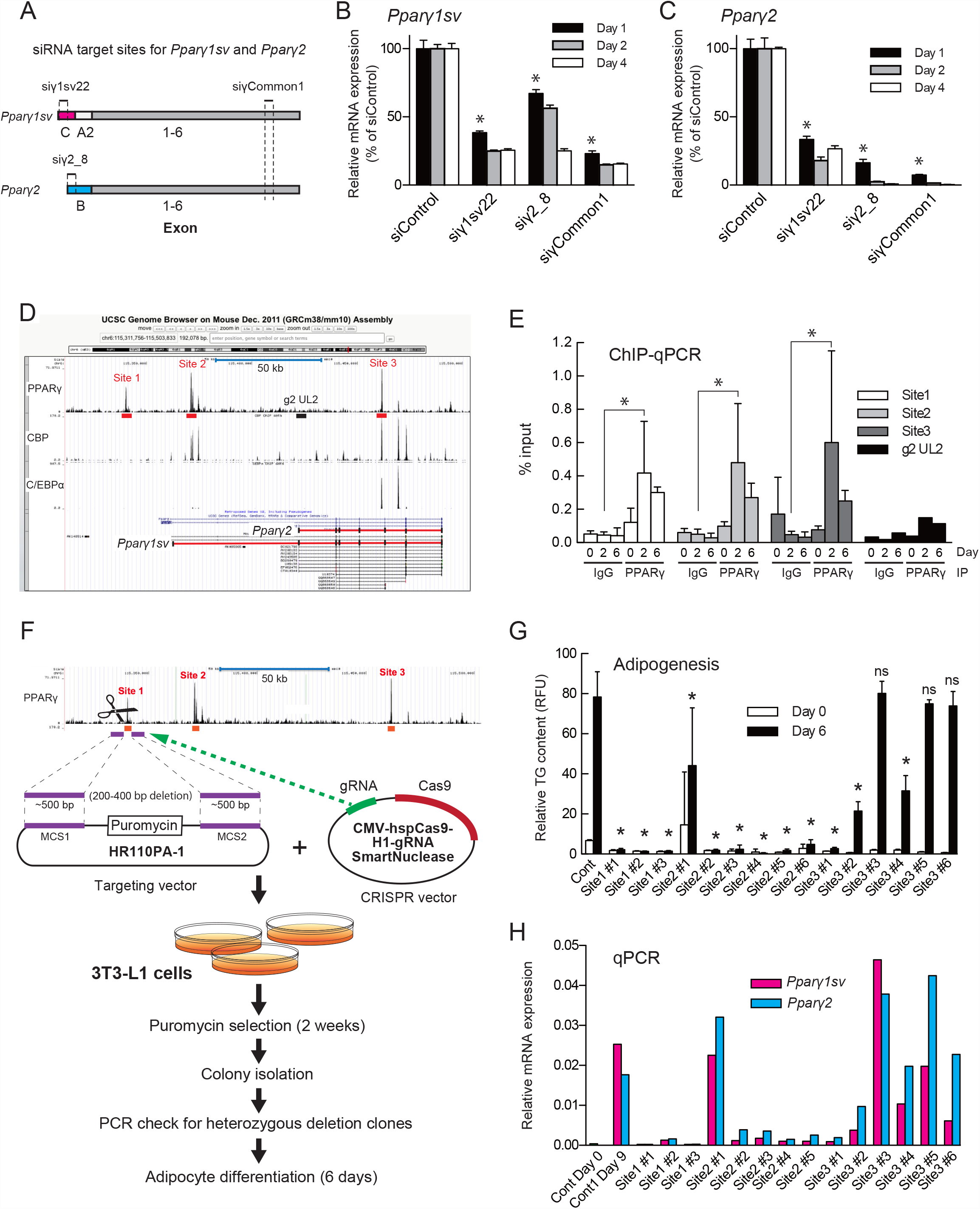
Binding of the PPARγ protein to the *Pparγ* gene locus regulates the expression and adipogenesis in 3T3-L1 cells: (A) Targeting sites of siRNAs against *Pparγ1sv* (siγ1sv22), *Pparγ2* (siγ2_8), and both transcripts (siγCommon1). Relative *Pparγ1sv* (B) and *Pparγ2* (C) mRNA levels in siRNA-treated 3T3-L1 cells on days 1, 2, and 4 of adipocyte differentiation determined by qPCR analysis. Values obtained for *Pparγ1sv* and *Pparγ2* were normalized to those of 18S rRNA. Values are shown as mean (SD) (*n* = 3). The asterisk (*) indicates *p* < 0.05 compared to siControl. (D) Binding sites of PPAR**γ**, CREB-Bindng Protein (CBP), and C/EBPα in the *Ppar****γ*** gene locus based on the densities of ChIP-seq data deposited in the NCBI (GSE20752). Sites 1– 3, candidate PPAR**γ**-binding sites; g2 UL2, non-PPAR**γ**-binding site. (E) Binding of PPARγ protein to candidate sites 1–3 in the *Ppar****γ*** gene locus was evaluated by ChIP-qPCR assay with anti-PPARγ antibody on days 0, 2, and 6 of adipocyte differentiation of 3T3-L1 cells. Values are shown as mean (SD) (*n* = 3). The asterisk (*) indicates *p* < 0.05 compared to negative controls with anti-IgG antibody. (F) Knockout (KO) strategy for PPAR**γ**-binding sites 1–3 using the CRISPR-Cas9 system with targeting vectors. (G) Relative triglyceride content in 3T3-L1 control and sites 1–3 KO clones on day 6 of adipocyte differentiation. Values are shown as mean (SD) (*n* = 4). The asterisk (*) indicates *p* < 0.05 compared to control cells. ns, not significant. (H) Relative *Pparγ1sv* and *Pparγ2* mRNA levels of the control and sites 1–3 KO 3T3-L1 on day 6 of adipocyte differentiation. Values were obtained by averaging the results of two independent experiments.

To further demonstrate the importance of these binding sites in adipocyte differentiation and the regulation of *Pparγ* expression, we established heterozygous knockout (KO) 3T3-L1 cells that lack each of three binding sites using the CRISPR-Cas9 system with the targeting vector (Fig. 1F). Upon adipocyte differentiation, the accumulation of intracellular triglyceride (TG) was largely hampered in site 1– and site 2–KO clones, but the KO effect was partial in site 3-KO clones (Fig. 1G), suggesting that sites 1 and 2 are indispensable for adipogenesis. Consistent with TG content, *Pparγ1sv* and *Pparγ2* mRNA levels were largely suppressed in site 1– and site 2–KO clones on day 9 of adipocyte differentiation, whereas those in site 3–KO clones were comparable with those of control cells (Fig. 1H).

Positive feedback mechanisms play vital regulatory roles in the expression of genes required for proper cellular differentiation, proliferation, and metabolic homeostasis (Ratushny et al., 2012). Although the presence of PPARγ-binding sites in the *Pparγ* gene locus (Mikkelsen et al., 2010; Waki et al., 2011) and the feedback regulation of *Pparγ* expression by PPARγ protein (Wakabayashi et al., 2009; Lee and Ge, 2014) in adipogenesis have been proposed previously, the studies basically focused on the *Pparγ2* expression, the major regulator of adipogenesis. In this study, we revealed that the specific knockdown of *Pparγ1sv* and *Pparγ2* resulted in the reciprocal downregulation of each transcript (Figs. 1B and 1C). Furthermore, we demonstrated that two of three PPARγ-binding sites in the *Pparγ* gene locus are critical elements for adipocyte differentiation (Fig. 1G) and *Pparγ* expression (Fig. 1H) in 3T3-L1 cells. Because these sites are shared between 3T3-L1 and mouse primary adipocytes (Siersbæk et al., 2012), we expect that the present findings can be applied to *in vivo* study for a more comprehensive understanding of the regulation of *Pparγ* expression during adipogenesis.

## Methods

### Cell culture and differentiation

The 3T3-L1 cells used in this study were obtained from JCRB Cell Bank (Osaka, Japan). Culture conditions, media, and method of adipocyte differentiation have been previously described (Takenaka et al., 2013).

### siRNA and qPCR

All siRNAs (Stealth RNAi) were purchased from Life Technologies (Carlsbad, CA, USA). Target sequences of *Pparγ1sv, Pparγ2*, and *Pparγ* common mRNAs were as follows: 5′-GAUCUGAAGGCUGCAGCGCUAAAUU-3′ (si*γ*1sv22), 5′-CCAGUGUGAAUUACAGCAAAUCUCU-3′ (si*γ*2_8), and 5′-CCAGGAGAUCUACAAGGACUUGUAU-3′ (si*γ*Common1). For siRNA transfection, 5 × 10^5^ 3T3-L1 cells per well were plated onto six-well plates and transfected using Lipofectamine RNAiMAX (Thermo Fisher Scientific, Waltham, MA, USA) according to the manufacturer’s instructions. To quantify *Pparγ1sv* and *Pparγ2* mRNA levels, total RNA was isolated from preadipocyte and adipocyte-differentiated 3T3-L1 cells using ISOGEN (Nippon Gene, Tokyo, Japan), reverse-transcribed using SuperScript III (Life Technologies), and amplified using Thunderbird SYBR qPCR mix (Toyobo, Osaka, Japan) in an ABI Prism 7900 HT sequence detection system (Life Technologies). Primer sequences for *Pparγ1sv, Pparγ2*, and *18S* ribosomal RNA (reference gene) qPCR analyses have been described elsewhere (Takenaka et al., 2013).

### ChIP-qPCR analyses

The 3T3-L1 cells cultured in 10-cm dishes were fixed with 1% formaldehyde in PBS at 25°C for 10 min, after which fixation was halted by addition of glycine solution. Immunoprecipitated protein-DNA complexes were prepared using the Magna ChIP A kit (Millipore, Burlington, MA, USA) according to the manufacturer’s instructions. Genomic DNA was immunoprecipitated with anti-PPARγ (A3409A, Perseus Proteomics, Tokyo, Japan) or anti-mouse IgG (Millipore, 12-371) antibody, purified, and amplified using Thunderbird SYBR qPCR mix (Toyobo) in an ABI Prism 7900 HT sequence detection system (Life Technologies) using the following site-specific primers: PPARg_ChIP1_UP2, 5′-GTACTTTTCTTTCTGGGTTTATTTTG-3′ and PPARg_Site1-5’_LP1, 5′-CTAATCTGGGTTAAGAGGATGTAATG-3′ for binding site 1; PPARg_ChIP_UP2, 5′-GGCTCAAAATACCCCTTCCATCTTA-3′ and PPARg_ChIP_LP2, 5′-GATGCTCAGGATTTGATGTCTCATA-3′ for binding site 2; PPARg_ChIP_UP3, 5′-CAGGTTTGATTCCCAACTCCCACATA-3′ and PPARg_ChIP_LP3, 5′-GATGATAGGCTACTTGTGAGCAAAGG-3′ for binding site 3; g2pro_ChIP_UP2, 5′-GCCTTTATTCTGTCAACTATTCCTTTT-3′ and g2pro_ChIP_LP2, 5′-AGTATTTATCTTTGGTTGAAACTCCTA-3′ for non-PPARγ-binding site.

### CRISPR/Cas9 genome editing

The CRISPR guide RNAs were cloned into the CMV-hspCas9-H1-gRNA SmartNuclease vector (System Biosciences, Palo Alto, CA, USA). The Cas9-target sequences for PPARγ-binding sites are as follows: 5′-CCTTCTAGCAGATCAAAAGT-3′ for site 1; 5′-TCAAAGAGTAAACCCACCAA-3′ for site 2; 5′-ACACTGTGATGCAGAGATGC-3′ for site 3. The upstream and downstream regions adjacent to the binding site were amplified using Advantage 2 polymerase (Takara Bio, Shiga, Japan) from 3T3-L1 genomic DNA and cloned into the HR110-PA-1 vector (System Biosciences). Primer sequences to amplify the 5′- and 3′-side genomic fragments for homologous recombination are as follows: PPARg_Site1-5’_UP1, 5′-CCTCGCTCCTGTTTTCTTTGTAAATA-3′ and PPARg_Site1-5’_LP1, 5′-CTAATCTGGGTTAAGAGGATGTAATG-3’ for 5′-side of site 1; PPARg_Site1-3′_UP1, 5′-CATGAAGAAGAAAGCAACCTCTGTATAA-3′ and PPARg_Site1-3′_LP1, 5′-CCTCCCCTCCCTTCTAATAAATGCTC-3′ for 3′-side of site 1; PPARg_Site2-5′_UP1, 5′-AGCTGGACTTTTGAGTTTTTCTCTAT-3′ and PPARg_Site2-5′_LP1, 5′-TTGAGCCACAAGACCAATATAAGATACA-3′ for 5′-side of site 2; PPARg_Site2-3′_UP1, 5′-TGCTAGGTAAACTGTTCGCCACTGAG-3′ and PPARg_Site2-3′_LP1, 5′-TGGTCGGGGTAAATCCTGTCTATG-3’ for 3′-side of site 2; PPARg_Site3-5′_UP1, 5′-CAATATGCTAAGTGCTGAGTGTAATGA-3′ and PPARg_Site3-5′_LP1, 5′-GTGCTCTTAACTGCTAAGCCATCTCTC-3′ for 5′-side of site 3; PPARg_Site3-3′_UP1, 5′-CCCACCATCATCTTGAGTGTCACATA-3′ and PPARg_Site3-3′_LP1, 5′-TTCCCCCAACCCTAAGCATTTCAACA-3′ for 3′-side of site 3. The Cas9 and targeting vectors were co-transfected into 3T3-L1 cells using PEI MAX™ (Cosmo Bio, Tokyo, Japan). For negative control clones, empty targeting vector was transfected into 3T3-L1 cells. After two days, the culture medium was replaced with DMEM containing 2.5 µg/mL puromycin, and the cells were cultured for two weeks. Single colonies were isolated using cell-cloning cylinders and expanded for an additional week. Heterozygous KO clones of the target site were screened by PCR.

### Intracellular TG content

Adipocyte differentiation was evaluated by staining cellular lipid droplets with the AdipoRed assay reagent (Lonza, Basel, Switzerland). The fluorescence intensity (excitation/emission = 485/535 nm) was measured using a Varioskan Flash (Thermo Fisher Scientific).

### Statistical analyses

We conducted one-way ANOVA followed by Tukey’s multiple comparison test. Differences between groups were considered significant at *p* < 0.05. All data were analyzed using GraphPad Prism 5.0 (GraphPad Software, San Diego, CA, USA) and are presented as the mean (SD) of the obtained values.

## Acknowledgments

We are grateful to Ms. Sawako Sato (Saitama Medical University) for her technical assistance. We would like to thank Editage (www.editage.jp) for English language editing.

## Funding

This study was supported by JSPS KAKENHI (https://www.jsps.go.jp/english/index.html) Grant number 25460299, 26461367, and 17K09866.

## Author Contributions

Yasuhiro Takenaka: conceptualization, data curation, investigation, methodology, visualization, and writing - original draft. Yoshihiko Kakinuma: conceptualization and supervision. Ikuo Inoue: conceptualization, supervision, resources, and writing - original draft.

## Notes

### Competing Interest Statement

The authors have declared no competing interest.

